# Developmental cold exposure increases the ability to maintain body temperature during future cold challenges in a free-living altricial bird

**DOI:** 10.64898/2026.09.24.754131

**Authors:** David A. Chang van Oordt, Conor C. Taff, Daniel R. Ardia, J. Ryan Shipley, Maren N. Vitousek

## Abstract

Developing in suboptimal temperatures can have widespread negative effects in organisms, but early exposure to thermal challenges can also trigger increased investment in thermoregulatory capacity. Within birds, the effects of cold exposure during incubation may differ between precocial and altricial species due to differences in the ontogeny of thermoregulation. To date, most of the evidence showing that embryonic cold exposure shapes future thermoregulatory capacity comes from a few precocial species in laboratory settings. Less is known about sensitivity to low-temperature incubation in altricial species which only become fully endothermic after hatching. In this study, we tested the effects of cold temperatures during embryonic development by temporarily cooling the nest boxes of free-living tree swallows (*Tachycineta bicolor*) during late incubation and subsequently exposing nestlings to an acute cold challenge. Cold exposure during early development made nestlings better at maintaining their body temperatures during the acute challenge. Developmentally cold-exposed birds also had higher thyroxine (T_4_) levels when sampled at lower ambient temperatures, whereas control birds showed the opposite relationship. Among nestlings that failed to maintain their body temperature during the acute cold challenge, developmentally cold-exposed nestlings secreted more corticosterone during the acute cold challenge than control nestlings. Despite these changes, cold-exposed birds did not differ from controls in cold-induced metabolic rates, body mass, or bacteria killing ability. These results are, to our knowledge, the first to show that developmental cold exposure increases the capacity to maintain body temperature during future cold challenges in a songbird. These findings suggest that experiencing even a mild decline in ambient temperatures during embryonic development could prime birds to cope more effectively with harsh or variable environmental conditions later in life.

## Introduction

It is well documented that experiences early in development can shape later-life phenotypes (Metcalfe & Monaghan, 2001; Nord & Giroud, 2020; Maccari et al., 2014; Weaver, 2009), but it is often unclear when and how these changes increase the capacity to cope with future challenges. On one hand, the “silver spoon hypothesis” posits that growing up in resource-rich environments with few challenges may allow organisms to develop robust phenotypes able to withstand challenging conditions, while challenging environments may disrupt of constrain development (Monaghan, 2007; Nettle & Bateson, 2015). On the other, developmental exposure to challenges may also promote phenotypes that can more easily cope with similar challenges in the future (hereafter called ‘priming’), although this flexibility may only be adaptive when developmental environments are good predictors of later-life environments (DeWitt et al., 1998; Monaghan, 2007; Mousseau & Fox, 1998; Nettle & Bateson, 2015). Whether a developmental challenge acts as a constraint or a cue may depend on its magnitude, with mild exposures priming compensatory development and severe exposures imposing costs. As we face a changing climate, understanding the role of environmental cues during development can help us understand how organisms cope with challenges and the causal agents of climate resilience (Fox et al., 2019; Pottier et al., 2022).

Climate change may increase variability in the conditions experienced by migratory birds, including exposure to unexpectedly cold weather during key life-history stages that shapes fitness and survival (Taff & Shipley, 2023). As temperatures rise, breeding birds can start nesting earlier in the breeding season, increasing the likelihood that both adults and offspring will have to withstand cold temperature events (Shipley et al., 2020). Unexpected temperatures during development can have profound and often detrimental effects on phenotypes, but there is also evidence in some species that early-life thermal conditions can improve the capacity to cope with future thermal challenges (Fallis et al., 2014; Gilchrist & Huey, 2001; Healy et al., 2019; *e.g.*, Pottier et al., 2022). In birds specifically, recent work has shown that developing in cold temperatures can have lasting effects (Nord & Giroud, 2020; Ruuskanen et al., 2021), but it is not clear if these effects prime individuals to cope with future periods of extreme cold. In precocial birds, incubation temperatures can modulate how juveniles respond to future thermal challenges (DuRant, Hopkins, & Hepp, 2011; Nichelmann, 2004; Shinder et al., 2002, 2011). For example, domestic turkeys (*Meleagris gallopavo*) exposed to colder incubation temperatures generate more heat later in life (Nichelmann, 2004), and wood ducks (*Aix sponsa*) incubated at cold temperatures later increase energy expenditure during a cold challenge, although this does not increase their ability to maintain higher body temperatures (DuRant et al., 2012). Thus, while there is some evidence of long-term phenotypic effects of incubation temperature in precocial birds, there is still little evidence that these changes translate to increased thermoregulatory capacity.

Few studies have evaluated the effects of incubation temperature on post-hatching phenotypes in altricial birds. The timing of development relative to hatching may determine when birds are most sensitive to varying thermal conditions (Starck & Ricklefs, 1998). In precocial birds, the hypothalamic-pituitary-adrenal (HPA) and the hypothalamic-pituitary-thyroid (HPT) axes of the endocrine system develop predominantly *in ovo*, and thus, the physiological stress response and machinery for endothermic homeostasis can be particularly sensitive to thermal inputs during this period (Debonne et al., 2008; Nichelmann, 2004; Nichelmann & Tzschentke, 2002). Altricial birds, in contrast, become endothermic after hatching, suggesting that thermal sensitivity may be greater during early nestling development (Dunn, 1975, 1979). Nord & Nilsson (2011) showed in blue tits (*Cyanistes caeruleus*) that incubation at low temperatures impaired growth but elevated basal metabolic rate. Similarly, a previous study in tree swallows *(Tachycineta bicolor)* found that nests that were experimentally cooled by an average of 5.9 °C produced smaller nestlings with reduced bactericidal ability, a measure of constitutive innate immunity (Ardia et al., 2010). Natural variation in exposure to cold during incubation also predicts stronger acute hormonal stress responses in adult tree swallows (Uehling et al., 2020).

We investigated whether ambient nest temperatures can shape the ability of tree swallow nestlings to cope with future thermal challenges. To our knowledge, no studies have directly evaluated the role of nest temperature on body temperature regulation in an altricial bird. Previous studies showing that cold exposure during incubation reduces nestling size suggest that thermoregulation may be impaired if nestlings are smaller (Ardia et al., 2010; Nord & Nilsson, 2011; Olson et al., 2008; Shipley et al., 2022). In blue tits, in particular, these size differences can persist into adulthood, potentially reducing their ability to respond to future cold events(Nord & Nilsson, 2011).

We also evaluated potential costs of low nest temperatures by assessing investment in the innate immune response. Exposure to low temperatures may constrain immune defence through energetic trade-offs, as resources allocated to meeting increased energetic demands may reduce those available for immune function (Iseri & Klasing, 2013). Such trade-offs are particularly likely when resources are scarce (French, DeNardo, et al., 2007) or when there energetic demands are greater (Albery et al., 2020; French, Johnston, et al., 2007; Nebel et al., 2012; Ruoss et al., 2019). Consequently, environmental conditions that require higher energy expenditure, like inclement weather, could limit available resources for physiological processes that are not immediately necessary and, thus, limit the strength of immune defences. There is ample evidence of proximate mechanisms that link ambient temperature and immune traits in birds (Hangalapura et al., 2004; Hegemann et al., 2012; Zhu et al., 2020), and a growing body of evidence shows that developmental environment can also shape long-term immune traits (Ardia et al., 2010; Burrows et al., 2019; DuRant, Hopkins, Hawley, et al., 2011; McDade, 2005). In particular, exposure to low incubation temperatures has been associated with reduced bactericidal ability in both tree swallows (Ardia et al., 2010) and Japanese quail (*Coturnix japonica*, Burrows et al., 2019).

Tree swallows may be especially vulnerable to cold conditions because declining ambient temperatures simultaneously increase thermoregulatory demands and reduce the availability of aerial insect prey, with severe cold events resulting in widespread nestling mortality (Shipley et al., 2020; Winkler et al., 2013). Additionally, females with experimentally cooled nests do not compensate by increasing the duration of incubation bouts; instead, they reduce incubation time and feed nestlings at lower rates (Ardia et al., 2010). The resulting nestlings are smaller in size and have reduced bacterial killing ability (Ardia et al., 2010).

It remains unclear, however, whether these phenotypic changes simply reflect the costs of developing in a suboptimal environment or reflect the adaptive prioritization of thermoregulatory development over investment in growth or immunity. Breeding early—when cold weather is more likely—is a strong predictor of reproductive success and fitness in tree swallows despite the apparent downsides of developing in cold conditions (Winkler et al., 2020). Nestlings whose parents breed earlier benefit from being fed insects with higher nutritional content because the availability of these insects typically peaks early in the breeding season (Twining et al., 2018). Thus, if exposure to cold during development enhances the capacity to cope with subsequent thermal challenges, these developmental responses could ultimately facilitate reproduction during the colder, but potentially advantageous, early part of the breeding season.

## Materials and Methods

### Study population and field methods

We studied tree swallow nestlings from a breeding population in Ithaca, NY (42°29′N, 76°27′W) between May and July 2022. The study site consists of two clusters of approximately 130 artificial nests made of wooden boxes set up in a grid. Within each cluster, boxes are ∼20 meters apart from each other (for a total of ∼250 available nesting sites). We monitor these nest boxes throughout the reproductive period, recording nesting activity and breeding stage following a standard protocol for tree swallows (see Winkler et al., 2020). All boxes are lined with an open corrugated cardboard box to allow easy removal of the nest to take measurements. We record the date when the first egg is laid and when the clutch is completed.

To measure the rates at which nestlings are fed by parents we installed a Radio Frequency Identification (RFID) receiver at each nest three days after clutch completion. Passive Integrated Transponder (PIT) tags attached to a colour band on the leg of each parent are detected and logged every time parents come to the hole of the nest box, which allows us to calculate nestling feeding rate using the protocol calibrated by our group (Vitousek et al., 2018). We captured each female six to eight days after their last egg was laid, and males six to twelve days after the eggs hatched (when possible); if birds were not yet banded, we fitted them with an individual metal band and PIT tag.

We also obtained hourly ambient temperature between April 1st and July 30th of 2022 from Cornell University’s Northeast Regional Climate Center (NRCC, https://www.nrcc.cornell.edu/) which has a weather station approximately 1.5 km from the field site.

### Experimental setup

Nest boxes were either assigned to an experimental cooling treatment (n = 26) or a control (n = 25). We artificially cooled the internal temperature of experimental nest boxes for five days during incubation following the methods by Vitousek et al. (Vitousek et al., 2022). In brief, we placed one ice pack (∼11 × 11 × 4 cm) under the cardboard liner separated by a film vapor barrier, and four ice packs in an ‘attic’ attached to the nest box the day before the treatment started. We ran this treatment on days 7–11 of the incubation period. To maintain the cooling effect, we swapped the thawed ice packs with frozen ones every four hours between 06:00 and 18:00. We prevented accumulated water condensation from dripping by placing paper towels around the sides of the nest box. Internal temperature in control nest boxes was allowed to vary naturally with ambient temperature. We measured internal box temperature by placing a temperature data logger (model MX2201, HOBO, Massachusetts, U.S.A.) on the wall of all experimental boxes, and of 13 control boxes. Neither incubation duration (Cold Mean ± S.E.: 13.9 ± 0.14, Control: 13.9 ± 0.13, *p* = 0.89), hatching date (Cold: June 1^st^ ± 0.47, Control: May 31^st^ ± 0.45, *p* = 0.42), hatching success (Cold: 0.48 ± 0.03, Control: 0.47 ± 0.03, *p* = 0.94) or nestling survival (Cold: 0.45 ± 0.03, Control: 0.44 ± 0.03, *p* = 0.92) differed between treatment groups demonstrating that embryonic mortality cannot account for the results of the treatments.

Nestlings were banded 12 days after they hatched. Upon removal from the nest, we weighed and collected a blood sample from all nestlings except for one randomly chosen nestling, which was left undisturbed in the nest until the next day when it was exposed to an experimental cold challenge. One blood sample from each nest was randomly selected for the bacteria killing assay, and the rest were pooled to assay thyroid hormones.

### Cold challenge and respirometry

Metabolic rates and body temperature regulation were measured on 13-day-old nestlings. Immediately upon removing the target nestling from the nest, we took a blood sample within the first three minutes after removal to measure circulating corticosterone (‘baseline’). Nestlings were then brought into the field lab where we weighed them, measured their pre-trial cloacal temperature using a high accuracy PT100 RTD input thermometer (Model HH804U, OMEGA Engineering Inc., Connecticut, U.S.A.), and placed them individually in a sealed chamber with temperature-controlled airflow. Airflow temperature was controlled by pumping air through copper tubing in an insulated chamber kept at 10 °C by a PELT 5 Peltier device (Sable Systems International—SSI—, Nevada, U.S.A.). We challenged one nestling from each nest for a total of 49 nestlings from 51 nests.

We measured the rate of oxygen consumption and carbon dioxide production during a 30-minute cold trial using a Foxbox Field Gas Analysis System (SSI). The air supply came from a pump sourcing natural air; thus, ambient oxygen concentration was set to 20.95% before each trial. Airflow was maintained at 360–440 mL min^-1^ and pumped through a sealed chamber either containing the nestling (animal chamber) or nothing (baseline chamber). Air from the chambers was diverted to an Intelligent Multiplexer (Model V3, SSI) set to pass air first from the control chamber for two minutes, then after flushing for four minutes, to switch to the animal chamber. Before the end of the cold trial and gas analysis, air was flushed again for four minutes and finally returned to the control chamber for two minutes. Airflow moved to an air humidity analyser (RH-300, SSI) and then dried using drierite (W. A. Hammond Drierite Company Ltd., Ohio, U.S.A.). After that, air moved to the CO_2_ analyser, was scrubbed with soda lime, and dried again before it moved to the O_2_ analyser in the Foxbox Field Gas Analysis System. At the end of the 30-minute cold challenge, we removed the nestling from the chamber, immediately recorded cloacal temperature, took a second blood sample for corticosterone analysis (‘cold-induced’), and returned it to its nest.

The resulting data files were processed using ExpeData version 1.9.27 (SSI) using a protocol recommended by the manufacturer. In brief, oxygen consumption (VO_2_) and carbon dioxide production (VCO_2_) were calculated from the difference between drift-corrected oxygen in the animal chamber and the baseline chamber using the following formulas:

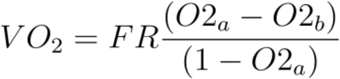

FR is the drift-corrected flow rate, O2 corresponds to the measured oxygen percentage, and the suffix “a” or “b” denotes whether the airflow comes from the animal or the baseline chamber, respectively. Finally, we took the mean of all VO_2_ estimates for a given animal as a measure of mean metabolic rate during the cold challenge.

### Sample processing

Upon collection, blood samples were kept on ice until they were processed less than four hours later. We separated plasma from red blood cells by spinning the samples at 3600 rpm for six minutes and removing it with a Hamilton syringe to be placed in a 0.6 mL microcentrifuge tube. We took 5 µL of the separated plasma for the bacteria killing assay, and the rest was stored in the field at -20 °C until it was transported to the lab where it was kept at – 80 °C until the thyroid hormone and corticosterone assays were done.

### Bacteria killing assay

We quantified BKA using a bacteria killing assay with blood plasma which measures activity of the complement system (Millet et al., 2007). By performing an assay that uses blood plasma, we can measure standing immune function of the complement system. We took 5 µL of plasma, diluted it to 2.5 and 1.25% in PBS and kept it at 4 °C until the start of the bacterial challenge. For the bacterial challenge, we followed the protocol used in Chang van Oordt et al. (2022) with some modifications. In brief, we prepared a stock suspension of *Escherichia coli* by reconstituting one pellet of Epower™ *E. coli* ATCC® 8739™ (Microbiologics, Minnesota, U.S.A.) in 40 µL of PBS prewarmed at 37 °C and letting it incubate in a water bath at the same temperature for 30 minutes. The stock solution was kept at 4 °C for no more than five days. On the day of the assay, we diluted 1 mL of the stock solution in 9 mL of PBS, to a final working concentration of 10^5^ colony forming units (CFU) mL^-1^. We then added 20 µL of working bacterial suspension to 100 µL of each plasma dilution in 0.6 mL tubes and incubated it at 41 °C in a water bath for 30 minutes. After incubation, we stopped the reaction by placing the tubes in a pre-chilled plastic tray for five minutes. After cooling, we plated 50 µL of each assay in two plates with tryptic soy agar, spread them throughout the agar with sterile glass beads, and let them incubate for 12–24 hours along with at least two positive controls that consisted of 50 µL of working solution. Finally, after incubation, we counted the number of CFUs in each plate assisted by CFU.Ai (mediXgraph Inc., California, U.S.A.).

We estimated BKA by dividing the number of bacterial colonies in each plate by the average number of colonies in all of each day’s positive control plates and subtracting from one. This represented the number of bacteria that were inhibited during the trial. This metric ranges from zero to one, but negative numbers are possible when bacterial growth rate is faster than bacterial inhibition during the trial (Tieleman et al., 2005).

### Plasma thyroid hormone concentrations

The HPT axis of the endocrine system regulates thermal homeostasis and energy expenditure. When body temperature drops, the thyroid gland secretes thyroxine (T_4_) in response to thyroid stimulating hormone (TSH) from the pituitary gland. T_4_ travels to different tissues where it is deiodinated into triiodothyronine (T_3_). T_3_ binding at nuclear and mitochondrial receptors alters gene expression, influencing thermoregulation, and often elevating metabolism (Darras et al., 2006; Zhang et al., 2018). We measured the average circulating T_3_ and T_4_ in blood plasma of 12-day-old nestlings. Because the minimum amount of plasma necessary for the assays exceeded the average amount of plasma that we can obtain from individual nestlings, we pooled plasma from multiple nestlings within the same nest. We had enough plasma for T_4_ measurements from 48 nests and 41 nests for T_3_. We sent each pooled plasma sample to the Animal Health Diagnostic Center at Cornell University to measure T_3_ in 75 µL of plasma using an IMMULITE® 2000 XPi system (Siemens Healthineers, Pennsylvania, U.S.A.) and T_4_ in 25 µL of plasma by radioimmunoassay.

### Circulating corticosterone concentration

We used a commercial corticosterone ELISA kit (DetectX Corticosterone Kit, K014-H5, Arbor Assays, Massachusetts, U.S.A.) that has been validated for tree swallow plasma corticosterone (Taff et al., 2019). We extracted steroids from blood plasma using a triple ethyl acetate extraction and measured corticosterone in duplicate using the kit. Extraction efficiency, calculated using corticosterone-spiked samples, was 91.93% and the mean intra-assay coefficient of variance was 10.80%.

### Statistical analyses

We ran all analyses using the program R (R Core Team, 2024). We ran a series of linear mixed effects and generalized linear mixed effects models with the functions *lmer* and *glmer* in the package *lme4* (Bates et al., 2015).

First, we compared the effect of treatment on the nestling feeding rate of female tree swallows using the best supported generalized linear mixed model reported by Vitousek et al. (Vitousek et al., 2018). We also included feeding rate as a potential predictor in the nestling linear models. To do this, we took the total number of feeding trips made by the female of a focal nest and divided it by the number of nestlings in her nest to obtain the mean total feeding rate for days 1–13 post-hatching. We labelled this the “mean feeding rate”.

We then used a model selection approach to test a series of linear mixed models using package *bbmle* (Bolker & R Development Core Team, 2020). All analyses consisted of comparing four main models for each response variable: one ‘full’ model containing treatment, ambient or cloacal temperature and a suite of potential confounding variables; one ‘temperature only’ model that included all predictors except for treatment group, one ‘interaction’ model that included all the previously mentioned predictors plus an interaction between treatment group and temperature; and one ‘reduced’ model that included all predictors except for temperature and treatment group. We used the Akaike Information Criterion for small samples (AICc) to evaluate the models. Within each model set we considered the best supported model to be the model with the lowest AICc scores; we also individually evaluated all additional models that were within a ΔAICc of two from the model with the lowest AICc score.

We tested the effects of treatment on body mass of 13-day-old nestlings using a full model that included date, the mother’s average feeding rate, and ambient temperature immediately preceding capture (defined here as the mean temperature during the last three hours prior to the trial). The analysis included 37 nestlings with complete data. We then tested the effects of the treatment on nestlings that received the secondary cold challenge using separate models for five response variables: initial cloacal temperature (immediately before the secondary cold challenge), final cloacal temperature (immediately after the secondary cold challenge), and mean rate of oxygen consumption during the secondary cold challenge, and baseline and cold-induced circulating corticosterone (before and after the challenge, respectively). The full model for initial cloacal temperature included date, brood size, body mass, ambient temperature immediately preceding the trial, and treatment group. Meanwhile, the full model for final cloacal temperature included date, body mass, duration of the trial, initial cloacal temperature, and treatment group. We also did a post hoc analysis of cloacal temperatures in which we compared the variance between treatments within a timepoint, and between timepoints within treatment using a two-tailed F-test for equality of two variances using the *var.test* function in the program R. For the full model of mean metabolic rate, we included body mass, initial body temperature, day of year, and treatment group. Lastly, the full model for baseline corticosterone included treatment, initial body temperature and day of year; while the full model for cold-induced corticosterone had treatment, final body temperature, time since nestling extraction, and baseline corticosterone, and day of year.

For nestlings that were did not go through the cold challenge, we built separate models to test the effects of treatment on circulating T_3_ concentration, circulating T_4_ concentration, and BKA in 12-day-old nestlings. For T_3_ and T_4_, we fit the full, temperature and interaction models with mean ambient temperature prior to sample collection, and we included a reduced model with no temperature predictor in it. The full model for T_3_ and T_4_ included date, mean nestling body mass, ambient temperature immediately preceding capture, and treatment group. In both thyroid hormone models we included ambient temperature immediately preceding capture (calculated as described above) because T_3_ peaks about two hours after cold exposure (Uribe et al., 1993). The T_3_ models additionally included circulating T_4_ concentration as a covariate. For BKA, we fit a full model that included day of year, body mass, mean ambient temperature 48 hours prior to sample collection, plasma dilution group, and number of CFUs in the positive control. For these analyses we included ambient temperature for 48 hours prior to sampling because we expect that BKA will be more strongly impacted by cold-induced changes in food availability than by cold itself. Low ambient temperatures reduce food supply in this system, like aerial insects (Winkler et al., 2013), and food limitation can cause shifts in immune investment via resource limitation and the stress response (Martin, 2009).

## Results

### Cold ambient nest temperatures prime nestling thermoregulation

We tested the hypothesis that early developmental cold exposure primes the development of thermoregulation in tree swallows, increasing subsequent thermoregulatory capacity. Specifically, we predicted that nestlings that hatched from eggs that were exposed to a five-day period of lower temperatures during incubation would later show elevated cold-induced metabolic rates and maintain higher body temperatures during exposure to cold. During the experimental cold exposure, the mean daytime temperature in cold-exposed nest boxes was lower by 1.98 °C than in control boxes (t = -3.23, df = 9.11, *p* = 0.01, Figure 2B), and the mean 24-hour temperature was lower by 1.78 °C (t = -3.38, *df* = 9.14, *p* = 0.008). Once the nestlings hatched, the parents compensated for colder conditions with more feeding visits, but only later in the nestling period (Figure 1). Peak feeding rates during the middle of the nestling provisioning period were higher among females whose nests were cold-treated during incubation (Figure 1). In the generalized linear mixed model of nestling feeding rate, there was a statistically significant interaction between nestling age squared and treatment group (ß = 1.03, 95% C.I. = [1.02, 1.05], *p* < 0.001, Supp. Table 12).

**Figure 1.**
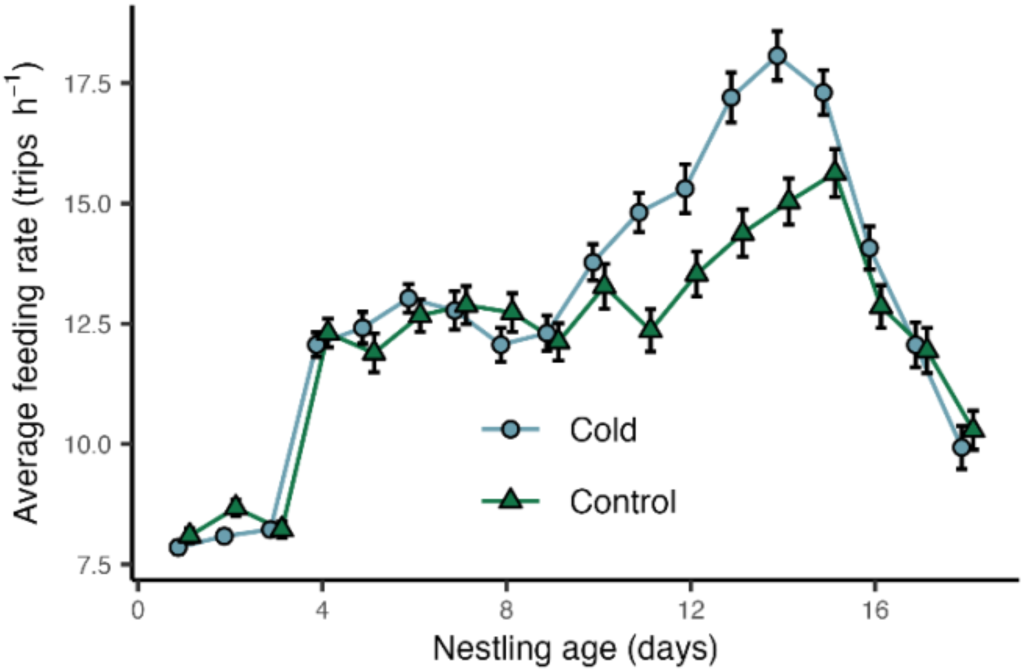
Females from cold-exposed nests feed nestlings more during the middle of nestling development than control females. Plot shows the mean feeding rate of female tree swallows by nestling age through the nestling period. Nestling age is shown as the number of days after hatching. Data is divided by treatment group. Points indicate the mean, and the error bar shows the standard error of the mean.

**Figure 2.**
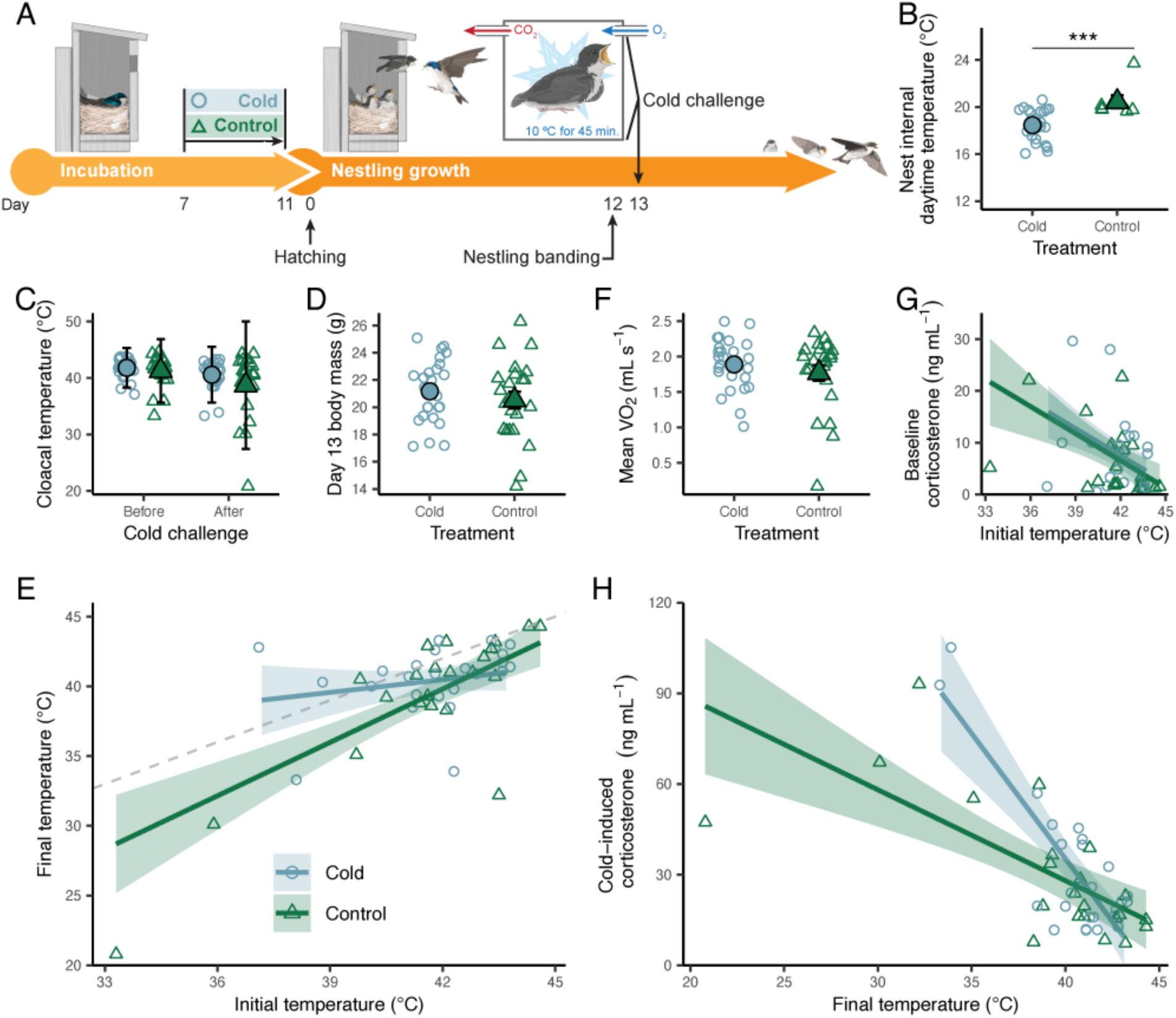
Cold-exposed nestlings (blue) lost less heat than control nestlings (green) when subjected to a cold challenge, and differed in their patterns of corticosterone secretion in response to the challenge. In plots **A**–**H**, filled shaped represent the means with their corresponding 95% confidence interval as an error bar, while open shapes represent individual data points. (**A**) Experimental design and timing of the cold challenge: cold exposure occurred during incubation, while the challenge occurred 13 days after hatching. (**B**) Mean and 95% C.I. daytime internal nest temperature. (**C**) Mean and 95% C.I. cloacal temperature of nestlings before and after the cold challenge. (**D**) Mean and 95% C.I. of 13-day-old nestling body mass. (**E**) Predicted temperature after the challenge as a function of the temperature at the start of the challenge for cold-exposed and control birds. The grey dashed line shows the point at which temperature does not change during the cold challenge. (**F**) Mean rate of oxygen consumption in cold-exposed and control nestlings. (**G**) Predicted baseline corticosterone as a function of cloacal temperature (initial, prior to the cold challenge) for cold-exposed and control nestlings based on the full regression model. (**H**) Predicted cold-induced circulating corticosterone concentration as a function of final cloacal temperature (post-challenge) for developmentally cold-exposed and control nestlings according to the best supported model.

To test for thermoregulatory capacity, we subjected 13-day-old nestlings to a cold challenge consisting of a chamber pumped with cold air (Figure 2A). Nestling body mass at 13 days old did not differ between developmentally cold-exposed and control nestlings (Figure 2D). The best supported model of body mass was the reduced model (Supp. Table 2–3), followed by the temperature model with a ΔAICc of 0.60 (Supp. Table 4). Date and mean feeding rate were positive predictors of body mass in both best supported models (Supp. Table 3–4). Ambient temperature was the only additional predictor in the temperature model, but within this model its effect was not statistically significant (Supp. Table 4).

Cold treatment did not affect body temperature prior to the cold challenge. In the best supported model for initial body temperature, higher ambient temperature immediately before capture predicted higher initial cloacal temperature (ß=0.28, CI = [0.14, 0.41], *p* < 0.001, Supp. Table S6), and nestlings that hatched earlier in the year had higher body temperatures (ß = - 0.31, CI = [-0.52, -0.09], p = 0.004, Supp. Table S6).

Meanwhile, cold-treated nestlings exhibited better thermoregulatory capacity by maintaining higher body temperatures during the challenge. The best supported model for final cloacal temperature included an interaction between treatment and initial temperature; no other models were within 5 ΔAICc (Supp. Table 7). In control birds, initial cloacal temperature positively predicted final cloacal temperature, but we did not see this relationship among developmentally cold-exposed nestlings (Figure 2E, Supp. Table 8). Instead, the final cloacal temperature of developmentally cold-exposed birds remained relatively high regardless of starting temperature (ß = 0.31, CI = [-0.21, 0.83], p = 0.247; Figure 2E; Supp. Table 8). The test of two variances showed that developmentally cold-exposed nestlings exhibited less cloacal temperature variance than did control nestlings before and after the cold trial (Initial: F = 0.39, Variance Ratio CI = [0.17, 0.88], p = 0.024; Final: F = 0.19, Variance Ratio CI = [0.08, 0.43], p < 0.001, Figure 2C). After the trial, variance increased in control but not in cold-exposed nestlings (Cold: F = 1.98, Ratio variance CI = [0.89, 4.42], p = 0.093; Control: F = 4.04, Ratio variance CI = [1.72, 9.54], p = 0.002; Figure 2C).

Despite increased thermoregulatory capacity, cold treatment during incubation did not affect the mean metabolic rate of nestlings during a 30-minute cooling trial. The best supported model of oxygen consumption rate during the cold challenge included initial cloacal temperature but not treatment (Figure 2F, Supp. Table 9), followed by the full model with 2.59 ΔAICc. In the temperature model, oxygen consumption increased with both body mass (ß = 0.09, CI = [0.06, 0.11], p < 0.001, Supp. Table 10) and initial cloacal temperature (ß = 0.07, [0.04, 0.10], p < 0.001, Supp. Table 10); and was lower in nestlings from nests that hatched earlier in the season (ß = 0.09, CI = [0.07, 0.12], *p* < 0.001, Supp. Table 10).

#### Lower ambient nest temperatures trigger higher sensitivity to the cold challenge

13-day-old nestlings showed elevated baseline corticosterone under colder ambient temperatures, but cold-treatment did not impact this relationship despite enhanced thermoregulatory capacity (Figure 2G). Among challenged nestlings, the best supported model of baseline corticosterone was the temperature model which included initial cloacal temperature and day of year (Supp. Table 11), which were both statistically significant predictors (Supp. Table 12). While baseline corticosterone was not impacted by treatment, developmental nest temperature did affect the regulation of corticosterone in response to an acute cold challenge. The best supported model of cold-induced corticosterone was the interaction model. Cold-induced corticosterone was higher the lower the nestling’s final temperature, but cold-exposed nestlings increased corticosterone secretion significantly more than control nestlings as their temperatures fell (Figure 2H, Supp. Figure 14).

### Developmental cold exposure alters the thermal responsiveness of circulating T_4_

Given that cold-treated nestlings were better at maintaining body temperature, we assessed HPT axis activity, which mediates thermoregulation (Debonne et al., 2008; Ruuskanen et al., 2021). We measured circulating levels of T_3_ and T_4_ in 12-day-old nestlings by pooling plasma samples by nest. Despite increased thermoregulatory capacity, cold-treated nestlings did not have higher baseline levels of circulating thyroid hormones (Figure 3A, B). However, we found T_4_ was more sensitive to ambient temperature in cold-treated nestlings than among controls (Figure 3C). The best supported model for circulating T_4_ included an interaction term between treatment and ambient temperature (Supp. Table 15–16). T_4_ in cold-exposed nestlings decreased with ambient temperature, but it increased among control nestlings (Figure 3C). Additionally, circulating T_4_ levels were higher in birds with greater body masses (ß = 0.07, CI = [0.01, 0.12], *p* = 0.015, Supp. Table 16). Meanwhile, developmental cold exposure did not affect circulating T_3_. The best supported model for T_3_ concentration was a reduced model (Supp. Table 17). T_3_ was positively predicted by mean body mass (ß = 5.93, CI = [3.11, 8.75], *p* < 0.001, Supp. Table 14) and circulating T_4_ concentration (ß = 16.28, CI = [2.25, 30.31], *p* = 0.024, Supp. Table 18).

**Figure 3.**
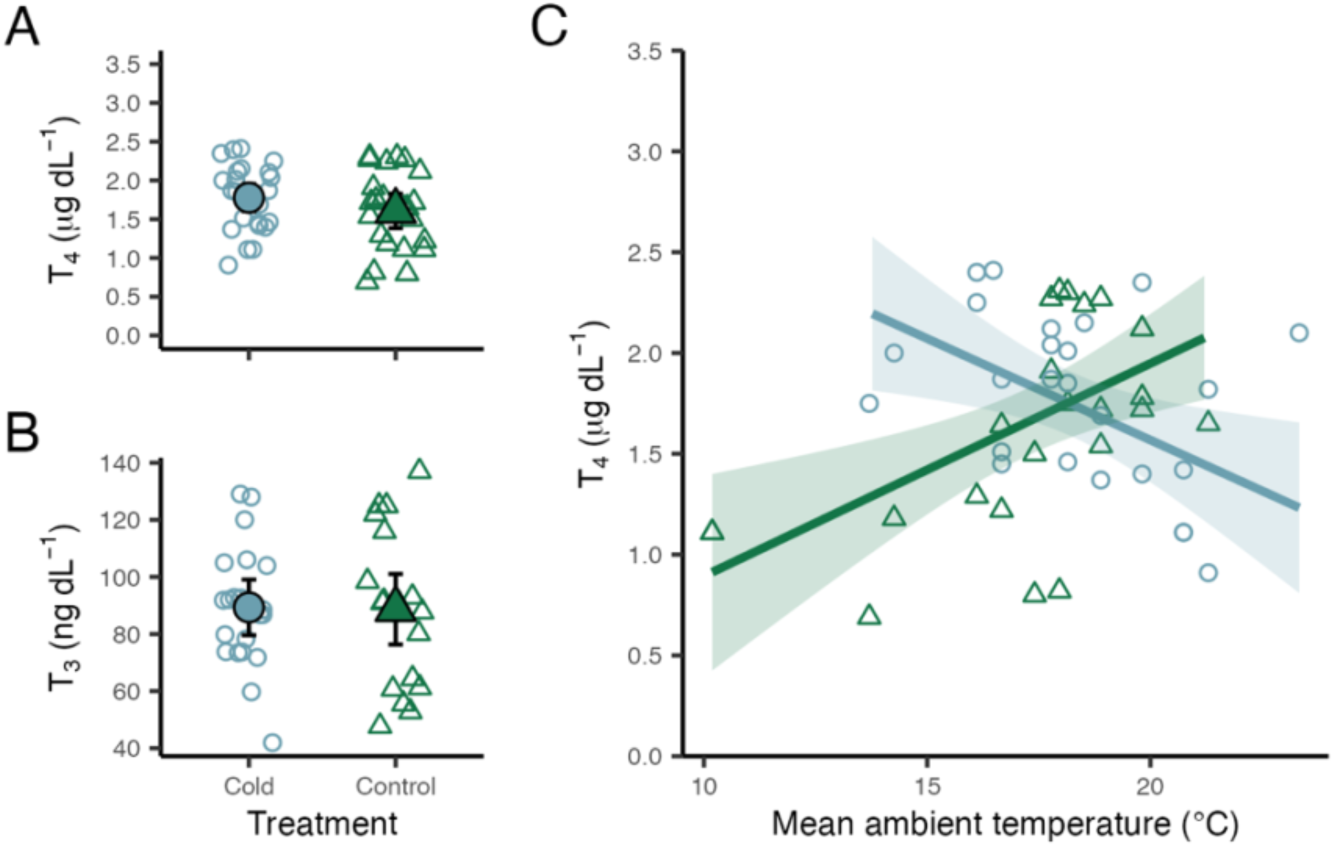
Thyroid hormone levels in 12-day-old nestlings. Mean of circulating T_4_ (**A**) and T_3_ (**B**) concentration. The filled points show the means divided by treatment group, and the error bars represent one standard error from the mean. In **C**, the lines show the predicted circulating T_4_ concentration as a function of ambient temperature immediately prior to sample collection for each treatment group as calculated by the interaction model. All other parameters in the model are set to the study population’s mean. In all cases, data are shown by the open points.

### Developmental cold exposure does not affect immune investment

We hypothesized that developmental cold exposure shifts resource allocation strategies to prioritize coping with inclement weather rather than investing in immune defences, traits that may be important over longer timescales. Thus, we assessed plasma bacteria killing ability (BKA), a measure of constitutive innate immune function, but we found no differences between treatment groups (Figure 4). Three models for BKA received similar support: the temperature model, the reduced model (ΔAICc = 0.16, Supp. Table 19), and the full model (ΔAICc = 0.46, Supp. Table 19). In all models, BKA was positively predicted by body mass, plasma dilution and CFU count in the positive control (Supp. Table 20–22).

**Figure 4.**
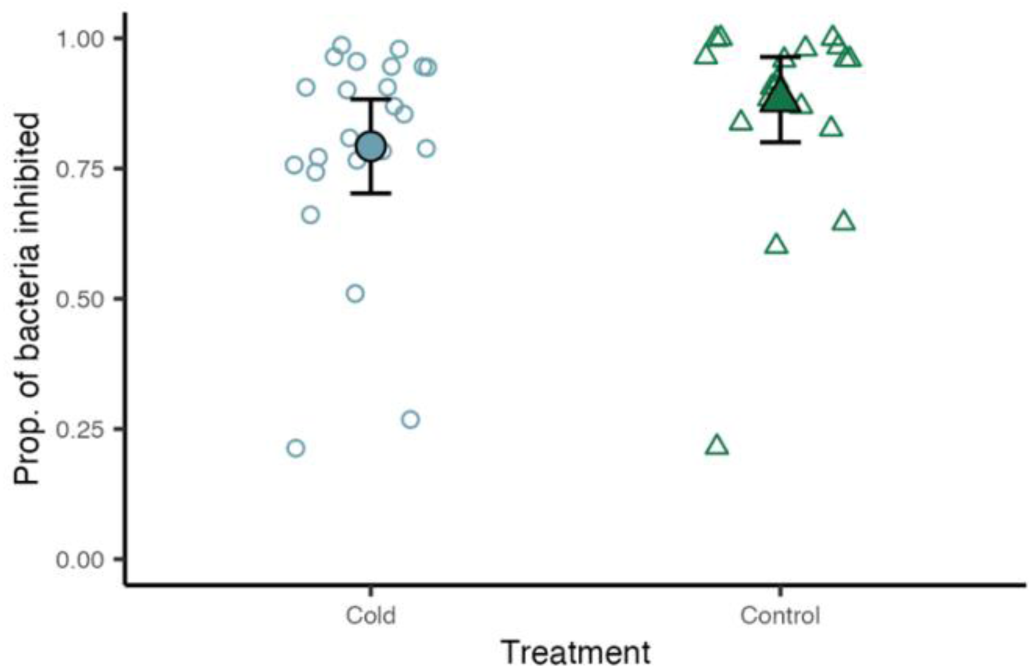
Mean and 95% confidence interval of the mean of the proportion of bacteria inhibited in the bacteria killing assay for each treatment group at 2.5% plasma dilution. Open circles and triangles show BKA value of each individual and the filled shapes represent the mean with one standard error represented in the error bar.

## Discussion

These results show that exposure to cooler temperatures during incubation increases the ability of nestling tree swallows to defend body temperature during a later cold challenge, and alters their glucocorticoid response to that challenge. In control nestlings, final cloacal temperature closely tracked initial temperature, so birds that began the challenge cooler also ended it cooler. In cold-exposed nestlings, final temperature was largely independent of initial temperature, and these birds were more likely than controls to finish the challenge near 41 °C, close to the normothermic body temperature of adult passerines (Prinzinger et al., 1991). Cold-exposed nestlings were also less variable in body temperature than controls before the challenge began, which indicates that differences in thermal stability were present at the start of the trial and were not produced solely by the challenge itself. Together, these results suggest that cold exposure during incubation prepares these altricial nestlings to cope with later cold conditions.

Few studies have tested whether the ability to maintain body temperature in cold conditions is affected by exposure to cold conditions early in development (during incubation) in birds. To our knowledge, all those that have tested this have done so in precocial species (*e.g.*, in wood ducks (DuRant et al., 2012), turkeys (Nichelmann, 2004), or chickens (Shinder et al., 2011)). In precocial species the endocrine mediators of thermoregulation—predominantly the HPT and HPA axes—develop prior to hatching, and thus, thermoregulatory capacity is expected to be particularly sensitive to thermal conditions during incubation in precocial birds. In contrast, in altricial birds, these axes mature over a longer time extending well into the post-hatching period. Thus, it is not clear whether the development of thermoregulatory capacity of altricial birds is similarly sensitive to temperature during incubation (Nord & Giroud, 2020; Ruuskanen et al., 2021). Our manipulation fell in the second half of incubation, when embryonic endocrine and thermoregulatory systems are differentiating but are not yet functional as they are after hatching. Our results indicate that the thermoregulatory ability of altricial species can be significantly impacted by even relatively small fluctuations in temperature during incubation (in this case a ∼2 °C reduction in nest temperature). Thus, we provide evidence that tree swallows could benefit from developmental exposure to cold temperatures that occur prior to hatching, if they subsequently face cold environmental conditions. To our knowledge, this is the first study in songbirds to show functional benefits of cold exposure during incubation. Because the timing of HPT and HPA maturation is far better described in precocial than in altricial embryos, future research should identify which components of these systems were sensitive to temperature during this window.

Short-term priming of the stress response has been previously reported in female adult tree swallows that experience cold temperatures after hatching (Vitousek et al., 2022). Meanwhile, tree swallow nestlings that experience cold temperatures as nestlings had increased sensitivity to stressors that persisted into adulthood suggesting long-term priming of the stress response (Uehling et al., 2020). Our data show that cold temperatures experienced as early as incubation are also followed by a stronger glucocorticoid response to a later cold challenge if nestling body temperature falls, suggesting that low temperatures during incubation may signal future adverse conditions. This result is notable because cold-exposed nestlings were less thermally challenged than controls during the trial: they cooled less yet secreted more corticosterone when their body temperature fell below 40 °C. A simple account in which better-prepared nestlings experience less stress would predict the opposite. One explanation is that the stronger glucocorticoid response is part of the mechanism of temperature defence rather than a separate outcome of treatment. Glucocorticoids mobilize the glucose and fatty acids that fuel thermogenesis, so a larger or faster response could support heat production during the early part of a cold exposure. Under this interpretation, improved thermoregulation and elevated corticosterone reflect one coordinated response to cold rather than two independently primed traits. Alternatively, a stronger glucocorticoid response could carry costs if it is activated repeatedly, and our data cannot distinguish between these possibilities. In birds, pre-natal exposure to corticosterone can dampen the stress response after hatching (Love & Williams, 2008; Zimmer et al., 2013), indicating that maternal stress could impact offspring phenotypes. Studies in mammals, however, suggest that mothers experiencing adversity during gestation can increase the sensitivity of the stress response of their offspring, although this pattern has not been consistent (McGowan & Matthews, 2018). Thus, it is unclear if low temperatures and maternal stress are similar signals of future adversity for the developing offspring. Birds deposit corticosterone in yolk during egg production (Hayward & Wingfield, 2004), which may only give a glimpse of the maternal experience with early adversity, thus limiting the value of the information received by the developing offspring. Temperature experienced during incubation, on the other hand, could be a better predictor of future conditions if the developing embryo is able to sense and adjust development accordingly. Altogether, our data show that pre-natal low temperatures can shape nestlings’ phenotypes, but it remains unclear how further experiences may calibrate this response during a bird’s lifespan.

Developmental cold exposure also altered the relationship between circulating T4 and ambient temperature. Nestlings that experienced cold temperatures during incubation had higher circulating levels of T_4_ when measured under lower ambient temperatures. Thyroid hormones play a critical role in the homeostatic control of body temperature (Zhang et al., 2018). Our results suggest that cold exposure may upregulate the HPT axis, preparing it to maintain body temperature when there is a high probability of facing inclement weather. Developmentally cold-exposed nestlings increased T_4_ as ambient temperature decreased, indicating activation of the physiological mechanisms for thermoregulation. Interestingly, however, control nestlings did not show this pattern, suggesting that development of the HPT axis is potentially accelerated in cold exposed nestlings compared to controls (Figure 3C).

Despite findings on T_4_, when controlling for circulating T_4_ we saw no effect of treatment on T_3_. T_3_ is the biologically active form of thyroid hormone which is enzymatically converted from T_4_ within the tissues. We measured T_3_ in plasma, but perhaps quantifying this hormone in more relevant tissues like pectoral muscles would show a different pattern. Taken together, these results suggest that elevated T_4_ in cold-exposed nestlings may act as a reservoir that allows for a faster thermoregulatory response when temperatures drop because cold-exposed nestlings may be able to produce more T_3_ out of a larger pool of precursor molecules. Work in other species has shown that cold exposure can increase the conversion of T_4_ to T_3_ (Collin et al., 2003; Van der Geyten et al., 1999); however, because we did not test thyroid hormone levels after the experimental cold challenge, we were not able to test that hypothesis here.

Surprisingly, our data suggest that cold-exposed nestlings maintained body temperature during the cold challenge without a detectable increase in mean oxygen consumption despite better thermoregulatory capacity in cold-exposed nestlings, although ambient temperature and body mass did (Supp. Table 10). As ambient temperatures decline below the optimum temperature range, the metabolic rates of homeotherms increase to generate heat (Cui et al., 2019; Swanson & Olmstead, 1999; Vézina et al., 2006; Williams & Tieleman, 2000). However, while elevating metabolic rate should allow individuals to maintain higher body temperatures during acute cold challenges, these processes do not always covary. For example, in wood ducks, developmental cold exposure increases cold-induced metabolic rates without influencing temperature regulation (DuRant et al., 2012, 2013). Two explanations for the patterns we report here are possible. First, cold-exposed nestlings may have retained heat more effectively, through feather insulation, fat stores, or peripheral vasoconstriction, none of which we measured. Second, the groups may have differed in the time course of heat production rather than in its average. A nestling that increases heat production quickly at the onset of cold and then reduces it once body temperature stabilizes could have the same mean oxygen consumption as a nestling that responds more slowly and continues to lose heat. Alternatively, increased non-shivering thermogenesis may have helped maintain body temperature while avoiding the high metabolic costs of shivering heat generation (Casagrande et al., 2020). However, non-shivering thermogenesis is not independent of cellular respiration, and our study did not find differences in metabolic rates or body mass. Distinguishing these explanations will require measuring the time course of oxygen consumption, or peak metabolic rate, together with body temperature. Despite the limitations explained above, however; our data suggest enhanced thermoregulatory efficiency in cold-exposed nestlings.

It is important to note that the finding that developmental cold exposure increases both T_4_ and the ability to maintain a constant body temperature during a cold challenge, but not cold-induced VO_2,_ may stem from high among-individual variation in measured VO_2_ due to uncontrolled environmental variation. Individual nestlings were sampled at different times of day; varying temperatures, stressors and times since last feeding, which can all affect metabolic rate, likely added additional variation to the dataset. It remains unclear if we would see the same results if we had been able to maintain stricter control of the nestling developmental environment. Moreover, in this study we only measured metabolic rates during the cold challenge because nestlings could only safely be removed from nests for up to ∼90 minutes; thus, we were not able to determine whether resting metabolic rates increased as a function of developmental cold exposure. However, this pattern has been suggested by studies in other species, including some altricial birds (Ben-Ezra & Burness, 2017; DuRant et al., 2012; DuRant, Hopkins, & Hepp, 2011; Nord & Nilsson, 2011; Olson et al., 2006).

With regards to immune investment, contrary to our predictions, cold exposure during development did not affect bacteria killing ability, a measure of innate immune function. These results differ from those of a previous study, also in tree swallows, that showed that nest cooling during late-stage incubation decreases BKA (Ardia et al., 2010). The most likely explanation is the magnitude of the manipulation. Nests in the earlier study were cooled by almost 6 °C, compared with approximately 2 °C here, and the stronger manipulation also reduced nestling body mass (Ardia et al., 2010), which we did not observe. Reduced nestling mass is a common result of lower incubation temperatures (*e.g.*, in zebra finches, *Taeniopygia guttata*; Olson et al., 2006, 2008; although not in blue tits Nord & Nilsson, 2011). Methodological differences may also contribute: we assayed BKA at 1.25% and 2.5% plasma dilutions, whereas Ardia et al. (2010) used 5%, and the lower concentrations may have increased sampling error in complement protein content. The pattern we observed could indicate that investment in thermogenic responses does not come at a cost to immune function, as has been seen in house sparrows (King & Swanson, 2013), or that the effects of cold exposure on immune function are not discernible until later in life, as in Japanese quail (Burrows et al., 2019). Taken together with Ardia et al. (2010), our results suggest that the consequences of cold exposure during incubation are dose dependent. A mild reduction in ambient nest temperature was followed by improved thermoregulation without detectable costs to growth or immunity, whereas a larger reduction imposed costs on both (although thermoregulatory capacity was not measured in that study). Whether developmental cold acts as a cue or as a constraint may therefore depend on how far temperatures fall, a possibility that could be tested by manipulating nest temperature across a range of magnitudes. Regardless, our results suggest that T_4_ and thermal regulation are more sensitive to small temperature decreases during incubation than body mass and BKA.

Overall, we found that early developmental experiences—as far back as *in ovo*—can impact a tree swallow’s phenotype after hatching. Specifically, exposure to a relatively mild decrease in temperature during embryonic development can prime nestlings to thermoregulate better and secrete more corticosterone as their body temperature drops during acute cold exposure, which can be beneficial for nestlings experiencing adverse weather. This study is, to our knowledge, the first to show thermoregulatory benefits of cold exposure during incubation and that pre-natal cold exposure alters the subsequent corticosterone response to cold in a songbird. Furthermore, we found that very small decreases (∼2 °C) in nest temperature during late incubation can have noticeable results in nestling phenotype. Cold-induced developmental plasticity may increase fitness, particularly in northern populations of tree swallows, by preparing nestlings for future inclement weather during key developmental stages as nestlings. By doing this study in near-natural conditions and in free-living birds that experience temperature fluctuations during incubation, we show that lower developmental temperatures can functionally benefit resilience to future cold events, at least during nestling growth. In other studies, incubation temperature is manipulated without fluctuations, so our study provides further evidence that these effects likely occur in nature.

Finally, in the era of climate change, thermal plasticity could provide an avenue for tolerating more variable or extreme environmental conditions (Calosi et al., 2007; Gunderson & Stillman, 2015; Khaliq et al., 2014; Pottier et al., 2022; Rodrigues & Beldade, 2020). We did not find evidence of a trade-off between thermoregulatory capacity and growth or constitutive innate immunity, suggesting that adaptation to more unpredictable environments may not necessarily impair immune function. However, it is unclear whether greater thermoregulatory investment comes at a cost to heat tolerance, other more energy-demanding arms of the immune response, downstream effects of elevated T_4_, or other traits. Future studies that investigate thermal sensitivity to different degrees of cold challenges, and to hyperthermic environments during development, will be important to further illuminate how thermoregulatory capacity is modulated, as well as the ability of tree swallows—and other altricial species—to withstand our changing climate.

## Acknowledgments

This project would not have been possible without the help of the 2022 tree swallow field crew: Gracey Brouilliard, Anthony Carnevale, Ava Ciaccia, David Jones, William Li, Audrey Su, Danielle Preston, Jennifer Houtz, Thomas Ryan, and Jennifer Uehling. This project was funded by the National Science Foundation BIO-IOS award no. 2128337 (CCT, MNV) and NSF-ORC award no. 2520505 (MNV) as well as the Richard H. and Mary Jane Schnoor Endowment Fund and the Athena Fund from the Cornell Lab of Ornithology (DCvO).

## Author contributions

DCvO, CCT, DRA and MNV conceived the ideas and designed the methodology, and the experiment was carried out and data collected by DCvO, CCT and DRA. Sample analysis was done by DCvO and CCT. DCvO did the data analyses and led the writing of the manuscript with substantial contributions from CCT, DRA, JRS and MNV. All authors contributed critically to the drafts and gave final approval for publication.

