## Supplementary Tables for "Developmental cold exposure increases the ability to maintain body temperature during future cold challenges in a free-living altricial bird"

Supplementary Materials

**Supplementary Table 1**

| ***Predictors*** | ***Incidence Rate Ratios*** | ***CI*** | ***p*** |
| --- | --- | --- | --- |
| (Intercept) | 14.64 | 13.27 – 16.15 | **<0.001***** |
| Temperature | 1.18 | 1.17 – 1.19 | **<0.001***** |
| Lay date | 0.97 | 0.92 – 1.03 | 0.317 |
| Brood size | 1.12 | 1.02 – 1.22 | **0.019*** |
| Brood sizeˆ2 | 1.03 | 0.97 – 1.09 | 0.280 |
| Nestling age | 1.04 | 1.02 – 1.05 | **<0.001***** |
| Nestling ageˆ2 | 0.77 | 0.76 – 0.78 | **<0.001***** |
| Hour | 1.02 | 1.02 – 1.03 | **<0.001***** |
| Hourˆ2 | 1.03 | 1.02 – 1.04 | **<0.001***** |
| Treatment [Control] | 0.90 | 0.77 – 1.06 | 0.207 |
| Nestling age* Treatment [Control] | 0.99 | 0.97 – 1.00 | 0.094 |
| Nestling age^2 * Treatment [Control] | 1.03 | 1.02 – 1.05 | **<0.001***** |
| Brood size * Treatment [Control] | 0.99 | 0.86 – 1.13 | 0.834 |
| Brood size^2 * Treatment [Control] | 0.99 | 0.88 – 1.12 | 0.861 |
| Brood size * Nestling age | 1.02 | 1.02 – 1.03 | **<0.001***** |
| **Random Effects** | | | |
| σ2 | 0.08 | | |
| τ00 | 0.02 | | |
| ICC | 0.24 | | |
| N | 38 | | |
| Observations | 9658 | | |
| Marginal R2 / Conditional R2 | 0.417 / 0.557 | | |
| **Supp. Table 1**. Model summary of the feeding rate generalized mixed model. All numerical variables were scaled to fit the model. | | | |

**Supplementary Table 2**

|  | **dAICc** | **df** | **weight** |
| --- | --- | --- | --- |
| Body mass ~ Date + Mean feeding rate | 0 | 4 | 0.50 |
| Body mass ~ Date + Mean feeding rate + Temperature (last 3h) | 0.60 | 5 | 0.37 |
| Body mass ~ Treatment + Date + Mean feeding rate + Temperature (last 3h) | 3.24 | 6 | 0.10 |
| Body mass ~ Treatment*Temperature (last 3h) + Date + Mean feeding rate | 5.91 | 7 | 0.03 |
| **Supp. Table 2**. Model selection table for the initial temperature models, described in the first column. | | | |

**Supplementary Table 3**

| ***Predictors*** | ***Estimates*** | ***CI*** | ***p*** |
| --- | --- | --- | --- |
| (Intercept) | -61.41 | -125.90 – 3.08 | 0.061 |
| Date | 0.48 | 0.09 – 0.87 | **0.018*** |
| Mean feeding rate | 0.01 | 0.00 – 0.01 | **0.001**** |
| Observations | 37 | | |
| R2 / R2 adjusted | 0.317 / 0.276 | | |
| **Supp. Table 3**. Summary of the reduced model of body mass showing the estimated predictor coefficients, their confidence intervals and the p-value. Statistically significant *p*-values are shown in bold. | | | |

**Supplementary Table 4**

| ***Predictors*** | ***Estimates*** | ***CI*** | ***p*** |
| --- | --- | --- | --- |
| (Intercept) | -53.09 | -117.96 – 11.79 | 0.105 |
| Date | 0.41 | 0.01 – 0.81 | **0.044*** |
| Mean feeding rate | 0.01 | 0.00 – 0.01 | **<0.001***** |
| Temperature (last 3h) | 0.14 | -0.07 – 0.36 | 0.176 |
| Observations | 37 | | |
| R2 / R2 adjusted | 0.354 / 0.295 | | |
| **Supp. Table 4**. Summary of the temperature model of body mass showing the coefficient estimates, their confidence intervals and *p*-values. Statistically significant *p*-values are shown in bold font. | | | |

**Supplementary Table 5**

|  | **dAICc** | **df** | **weight** |
| --- | --- | --- | --- |
| Cloacal temp. (initial) ~ Brood Size + Body mass + Temperature (last 3h) + Date | 0 | 6 | 0.75 |
| Cloacal temp. (initial) ~ Treatment + Brood Size + Body mass + Temperature (last 3h) + Date | 2.69 | 7 | 0.20 |
| Cloacal temp. (initial) ~ Treatment*Temperature (last 3h) + Brood Size + Body mass + Date | 5.49 | 8 | 0.05 |
| Cloacal temp. (initial) ~ Brood Size + Body mass + Date | 13.80 | 5 | 0.00 |
| **Supp. Table 5**. Model descriptions and AICc scores of the initial cloacal temperature models. The table includes the ΔAICc, degrees of freedom and model weight. | | | |

**Supplementary Table 6**

| ***Predictors*** | ***Estimates*** | ***CI*** | ***p*** |
| --- | --- | --- | --- |
| (Intercept) | 79.77 | 45.64 – 113.91 | **<0.001***** |
| Brood Size | 0.02 | -0.46 – 0.50 | 0.934 |
| Body mass | 0.32 | 0.11 – 0.53 | **0.004**** |
| Temperature (last 3h) | 0.28 | 0.14 – 0.41 | **<0.001***** |
| Date | -0.31 | -0.52 – -0.09 | **0.01**** |
| Observations | 51 | | |
| R2 / R2 adjusted | 0.479 / 0.434 | | |
| **Supp. Table 5**. Summary of the temperature model of initial cloacal temperature. The table includes the coefficient estimates, their confidence intervals, and p-values. Statistically significant *p*-values are shown in bold font. | | | |

**Supplementary Table 7**

|  | **dAICc** | **df** | **weight** |
| --- | --- | --- | --- |
| Cloacal temp. (final) ~ Treatment*Cloacal temp. (initial) + Body mass + Trial duration + Date | 0 | 8 | 0.91 |
| Cloacal temp. (final) ~ Treatment + Body mass + Trial duration + Cloacal temp. (initial) + Date | 5.93 | 7 | 0.05 |
| Cloacal temp. (final) ~ Body mass + Trial duration + Cloacal temp. (initial) + Date | 5.96 | 6 | 0.05 |
| Cloacal temp. (final) ~ Body mass + Trial duration + Date | 25.26 | 5 | 0.00 |
| **Supp. Table 7**. Model description and selection criteria for final cloacal temperature models. The table shows the ∆AICc, degrees of freedom and corresponding weight for each model. | | | |

**Supplementary Table 8**

| ***Predictors*** | ***Estimates*** | ***CI*** | ***p*** |
| --- | --- | --- | --- |
| (Intercept) | 3.90 | -55.44 – 63.25 | 0.895 |
| Treatment [Control] | -41.44 | -69.78 – -13.09 | **0.005**** |
| Cloacal temp. (initial) | 0.31 | -0.22 – 0.83 | 0.247 |
| Body mass | 0.64 | 0.35 – 0.93 | **<0.001***** |
| Trial duration | 0.00 | -0.00 – 0.01 | 0.267 |
| Date | -0.02 | -0.32 – 0.28 | 0.881 |
| Treatment [Control] * Cloacal temp. (initial) | 0.97 | 0.29 – 1.65 | **0.006**** |
| Observations | 49 | | |
| R2 / R2 adjusted | 0.764 / 0.730 | | |
| **Supp. Table 8**. Summary of the interaction model of cloacal temperature after the cold challenge. The table shows the coefficient estimates, their confidence intervals and their *p*-values. Statistically significant *p*-values are shown in bold font. | | | |

**Supplementary Table 9**

|  | **dAICc** | **df** | **weight** |
| --- | --- | --- | --- |
| VO2 ~ Body mass + Cloacal temp. (initial) + Date | 0.00 | 5 | 0.74 |
| VO2 ~ Treatment + Body mass + Cloacal temp. (initial) + Date | 2.59 | 6 | 0.20 |
| VO2 ~ Treatment*Cloacal temp. (initial) + Body mass + Date | 5.10 | 7 | 0.06 |
| VO2 ~ Body mass + Date | 17.53 | 4 | 0.00 |
| **Supp. Table 9**. Model definitions and selection criteria for the oxygen consumption models. The table includes ∆AICc, degrees of freedom and weight of each model. | | | |

**Supplementary Table 10**

| ***Predictors*** | ***Estimates*** | ***CI*** | ***p*** |
| --- | --- | --- | --- |
| (Intercept) | -18.14 | -22.76 – -13.52 | **<0.001***** |
| Body mass | 0.09 | 0.06 – 0.11 | **<0.001***** |
| Cloacal temp. (initial) | 0.07 | 0.04 – 0.10 | **<0.001***** |
| Date | 0.09 | 0.07 – 0.12 | **<0.001***** |
| Observations | 49 | | |
| R2 / R2 adjusted | 0.800 / 0.787 | | |
| **Supp. Table 10**. Summary of the temperature model of oxygen consumption during the cold challenge. The table includes the coefficient estimates, their confidence intervals and *p*-values. Statistically significant *p*-values are shown in bold font. | | | |

**Supplementary Table 11**

|  | **dAICc** | **df** | **weight** |
| --- | --- | --- | --- |
| Baseline CORT ~ Cloacal temp. (initial) + Date | 0.00 | 4 | 0.57 |
| Baseline CORT ~ Treatment + Cloacal temp. (initial) + Date | 2.33 | 5 | 0.18 |
| Baseline CORT ~ Date | 2.60 | 3 | 0.15 |
| Baseline CORT ~ Treatment*Cloacal temp. (initial) + Date | 3.48 | 6 | 0.10 |
| **Supp. Table 11**. Model definitions and selection criteria for the baseline corticosterone models. The table includes ∆AICc, degrees of freedom and weight of each model. | | | |

**Supplementary Table 12**

| ***Predictors*** | ***Estimates*** | ***CI*** | ***p*** |
| --- | --- | --- | --- |
| (Intercept) | -242.09 | -361.36 – -122.81 | **<0.001***** |
| Cloacal temp. (initial) | -0.78 | -1.48 – -0.07 | **0.032*** |
| Date | 1.72 | 1.05 – 2.39 | **<0.001***** |
| Observations | 49 | | |
| R^2^ / R^2^ adjusted | 0.458 / 0.435 | | |
| **Supp. Table 12**. Summary of the temperature model of baseline corticosterone prior to the cold challenge. The table includes the coefficient estimates, their confidence intervals and p-values. Statistically significant p-values are shown in bold font. | | | |

**Supplementary Table 13**

|  | **dAICc** | **df** | **weight** |
| --- | --- | --- | --- |
| Stress-induced Corticosterone ~ Treatment * Cloacal temp. (final) + Time since first disturbance + Baseline corticosterone + Date | 0.00 | 8 | 0.99 |
| Stress-induced Corticosterone ~ Cloacal temp. (final) + Time since first disturbance + Baseline corticosterone + Date | 10.09 | 6 | 0.01 |
| Stress-induced Corticosterone ~ Treatment + Cloacal temp. (final) + Time since first disturbance + Baseline corticosterone + Date | 11.24 | 7 | 0.00 |
| Stress-induced Corticosterone ~ Time since first disturbance + Baseline corticosterone + Date | 35.48 | 5 | 0.00 |
| **Supp. Table 13**. Model definitions and selection criteria for the stress-induced corticosterone models. The table includes ∆AICc, degrees of freedom and weight of each model. | | | |

**Supplementary Table 14**

| ***Predictors*** | ***Estimates*** | ***CI*** | ***p*** |
| --- | --- | --- | --- |
| (Intercept) | 479.54 | 51.50 – 907.58 | **0.029*** |
| Treatment [Control] | -218.57 | -333.31 – -103.83 | **<0.001***** |
| Cloacal temp. (final) | -8.29 | -10.92 – -5.66 | **<0.001***** |
| Time since first disturbance | -0.31 | -1.34 – 0.73 | 0.555 |
| Baseline corticosterone | -0.28 | -1.15 – 0.59 | 0.519 |
| Date | -0.55 | -3.05 – 1.95 | 0.66 |
| Treatment [Control] * Cloacal temp. (final) | 5.28 | 2.44 – 8.13 | **0.001**** |
| Observations | 49 | | |
| R^2^ / R^2^ adjusted | 0.614 / 0.559 | | |
| **Supp. Table 14**. Summary of the interaction model of stress-induced corticosterone during the cold challenge. The table includes the coefficient estimates, their confidence intervals and p-values. Statistically significant p-values are shown in bold font. | | | |

**Supplementary Table 15**

|  | **dAICc** | **df** | **weight** |
| --- | --- | --- | --- |
| T_4_ ~ Treatment*Temperature (last 3h) + Date + Body mass | 0 | 7 | 0.96 |
| T_4_ ~ Date + Body mass | 7.04 | 4 | 0.03 |
| T_4_ ~ Date + Body mass + Temperature (last 3h) | 8.90 | 5 | 0.01 |
| T_4_ ~ Treatment + Date + Body mass + Temperature (last 3h) | 11.16 | 6 | 0.00 |
| **Supp. Table 15**. Model definitions and selection criteria for circulating T_4_ concentration models. The table shows the ∆AICc, degrees of freedom and weights of each model. | | | |

**Supplementary Table 16**

| ***Predictors*** | ***Estimates*** | ***CI*** | ***p*** |
| --- | --- | --- | --- |
| (Intercept) | -3.28 | -12.02 – 5.45 | 0.452 |
| Treatment [Control] | -3.74 | -5.73 – -1.76 | **<0.001***** |
| Temperature (last 3h) | -0.10 | -0.18 – -0.02 | **0.014*** |
| Date | 0.03 | -0.02 – 0.09 | 0.223 |
| Body mass | 0.06 | 0.00 – 0.12 | **0.036*** |
| Treatment [Control] * Temperature (last 3h) | 0.21 | 0.10 – 0.32 | **<0.001***** |
| Observations | 48 | | |
| R2 / R2 adjusted | 0.357 / 0.281 | | |
| **Supp. Table 12**. Model summary of the interaction model of circulating T_4_ concentration. The table shows coefficient estimates, their confidence intervals and their *p*-values. Statistically significant *p*-values are shown in bold font. | | | |

**Supplementary Table 17**

|  | **dAICc** | **df** | **weight** |
| --- | --- | --- | --- |
| T_3_ ~ Date + Body mass + T4 | 0 | 5 | 0.70 |
| T_3_ ~ Date + Body mass + Temperature (last 3h) + T4 | 2.76 | 6 | 0.18 |
| T_3_ ~ Treatment + Date + Body mass + Temperature (last 3h) + T4 | 4.43 | 7 | 0.08 |
| T_3_ ~ Treatment*Temperature (last 3h) + Date + Body mass + T4 | 5.63 | 8 | 0.04 |
| **Supp. Table 13**. Model definitions and selection criteria for the circulating T_3_ concentration models showing the ∆AICc values, degrees of freedom and model weight. | | | |

**Supplementary Table 18**

| ***Predictors*** | ***Estimates*** | ***CI*** | ***p*** |
| --- | --- | --- | --- |
| (Intercept) | 217.80 | -160.96 – 596.57 | 0.251 |
| Date | -1.69 | -3.99 – 0.61 | 0.146 |
| Body mass | 5.93 | 3.11 – 8.75 | **<0.001***** |
| T_4_ | 16.28 | 2.25 – 30.31 | **0.024*** |
| Observations | 41 | | |
| R2 / R2 adjusted | 0.460 / 0.416 | | |
| **Supp. Table 14**. Model summary of the reduced model for circulating T_3_ concentration. The table shows the coefficient estimates, their confidence intervals and *p*-values. | | | |

**Supplementary Table 19**

|  | **dAICc** | **df** | **weight** |
| --- | --- | --- | --- |
| BKA ~ Date + Body mass + Mean ambient temperature (last 48h) + Plasma dilution + CFU count in positive control | 0.00 | 7 | 0.34 |
| BKA ~ Date + Body mass + Plasma dilution + CFU count in positive control | 0.16 | 6 | 0.31 |
| BKA ~ Treatment + Date + Body mass + Mean ambient temperature (last 48h) + Plasma dilution + CFU count in positive control | 0.46 | 8 | 0.27 |
| BKA ~ Treatment*Mean ambient temperature (last 48h) + Date + Body mass + Plasma dilution + CFU count in positive control | 2.91 | 9 | 0.08 |
| Supp. Table 19. Model definitions and selection criteria for the BKA models. The table includes ∆AICc, degrees of freedom and weight of each model. | | | |

**Supplementary Table 20**

| ***Predictors*** | ***Estimates*** | ***CI*** | ***p*** |
| --- | --- | --- | --- |
| (Intercept) | 2.28 | -1.82 – 6.38 | 0.272 |
| Date | -0.02 | -0.05 – 0.01 | 0.172 |
| Body mass | 0.02 | 0.00 – 0.04 | **0.014*** |
| Temperature (last 48h) | 0.03 | -0.01 – 0.07 | 0.126 |
| Plasma dilution | 30.04 | 23.59 – 36.49 | **<0.001***** |
| Positive CFU count | -0.00 | -0.00 – -0.00 | **<0.001***** |
| Observations | 89 | | |
| R2 / R2 adjusted | 0.597 / 0.573 | | |
| **Supp. Table 20**. Summary of the temperature model of BKA. The table includes the coefficient estimates, their confidence intervals and p-values. Statistically significant p-values are shown in bold font. | | | |

**Supplementary Table 21**

| ***Predictors*** | ***Estimates*** | ***CI*** | ***p*** |
| --- | --- | --- | --- |
| (Intercept) | 0.24 | -2.93 – 3.40 | 0.883 |
| Date | -0.00 | -0.02 – 0.02 | 0.687 |
| Body mass | 0.03 | 0.01 – 0.04 | **<0.001***** |
| Plasma dilution | 30.15 | 23.65 – 36.65 | **<0.001***** |
| Positive CFU count | -0.00 | -0.00 – -0.00 | **<0.001***** |
| Observations | 89 | | |
| R2 / R2 adjusted | 0.586 / 0.566 | | |
| **Supp. Table 21.** Summary of the reduced model of BKA. The table includes the coefficient estimates, their confidence intervals and *p*-values. Statistically significant *p*-values are shown in bold font. | | | |

**Supplementary Table 22**

| ***Predictors*** | ***Estimates*** | ***CI*** | ***p*** |
| --- | --- | --- | --- |
| (Intercept) | 2.20 | -1.88 – 6.28 | 0.287 |
| Treatment [Control] | 0.05 | -0.03 – 0.14 | 0.181 |
| Date | -0.02 | -0.05 – 0.01 | 0.177 |
| Body mass | 0.02 | 0.00 – 0.04 | **0.016*** |
| Temperature (last 48h) | 0.03 | -0.01 – 0.07 | 0.112 |
| Plasma dilution | 29.89 | 23.46 – 36.31 | **<0.001***** |
| Positive CFU count | -0.00 | -0.00 – -0.00 | **<0.001***** |
| Observations | 89 | | |
| R2 / R2 adjusted | 0.606 / 0.577 | | |
| **Supp. Table 22**. Summary of the full model of BKA. The table includes the coefficient estimates, their confidence intervals and p-values. Statistically significant p-values are shown in bold font. | | | |
